# Adaptive potential and evolutionary divergence of scototaxis in juvenile Trinidadian Guppies

**DOI:** 10.64898/2026.05.01.722148

**Authors:** Emily Phelps, Lengxob Yong, Pamela Prentice, Bonnie Fraser, Erik Postma, Alastair Wilson

**Affiliations:** Centre for Ecology and Conservation, University of Exeter, Cornwall UK; Department of Biology, Davidson College, Davidson, North Carolina US; Department of Veterinary Bioscience, SRUC, Edinburgh, UK; Department of Biosciences, University of Exeter, Exeter UK

**Author notes:** **Ethics statement** Behavioural experiments were conducted under licence from the Home Office (UK) and under the auspices of the Animal (Scientific Procedures) 1986 Act (PPL 30/3256) and with local ethical approval from the University of Exeter.

**Keywords:** boldness, habitat choice, predation, behavioural variation, local adaptation

## Abstract

Habitat choice and resource acquisition represent an intrinsic trade-off, resulting in variation among individuals and potentially local adaptation among populations. Here, we quantify scototaxis, or preference for dark versus light backgrounds, in juvenile guppies. Scototaxis assays, in which individuals choose between safer dark and riskier light backgrounds, are widely used as a measure of boldness in small fishes, but they can equally be interpreted as choice between two habitats. By rearing and testing 586 fish descended from ten natural populations from Trinidad under common garden conditions, we first quantify (broad sense) heritable variation, i.e. evolutionary potential, within populations. Next, we test for evolutionary divergence among populations in mean preference, and if present, whether ancestral predation regime is a mediator of divergence. Finally, we ask whether families and/or populations differ in the amount of behavioural variation they contain. Preference varied among families (12% of total variance), consistent with broad-sense heritable variation. We also found that mean preference varied among populations (11% of total variance explained). Evolutionary divergence among-populations is partly explained by ancestral predation regime, with populations from low-predation sites showing a stronger preference for dark backgrounds than high-predation populations from the same river. Additionally, we find that within-population behavioural variation is greater in high-predation populations. We conclude that guppy populations contain heritable variation that could facilitate adaptive evolution if scototaxis is subject to natural selection. Furthermore, while genetic drift may also contribute to evolutionary divergence among-populations, observed patterns are qualitatively consistent with local adaption to predation regime. Our results suggests that high predation sites favour ‘bolder’ habitat choice on average, but also that local predation regime shapes the evolutionary dynamics of variation, perhaps by maintaining shy-bold variation among individuals, or by favouring individuals with less-predictable behaviour.

## Introduction

Gaining resources whilst evading predation represents a daily plight for many animals in the wild. Subsequently, habitat choice and boldness, the behavioural response to perceived risk, are functionally related because the former often depends on a trade-off between risk (e.g. exposure to predators) and reward (e.g. access to food resources or mating opportunities; Milinski & Heller, 1978; McNamara & Houston, 1987). Bolder (i.e. more risk prone) individuals will be more likely to use high-risk habitats when foraging, potentially accessing additional, higher-quality, or less-exploited resources (Brown & Kotler, 2004; Dammhahn & Almeling, 2012; Steinhoff *et al*., 2020; Erixon *et al*., 2025). Individual variation in habitat choice, and boldness, reflects how each individual solves this trade-off, which may also be influenced by differential predation risk, physiological state and competition (Brown & Kotler, 2004; Sih & Del Giudice, 2012). While some habitats may be universally preferable over others because of high resources and/or low risk, habitat preference may also vary among individuals within populations (Camacho *et al*., 2020). When preference is linked to other traits, this leads to a non-random distribution of phenotypes in the environment (Edelaar and Bolnick, 2019). For example, sticklebacks originating from a lake environment are more likely to migrate into stream environments if they already possess a ‘stream-like’ morphology (Bolnick *et al*., 2009). In this example, individual habitat choice is adaptive and dependent on a phenotype (morphology) that is known to be heritable. From this, it follows that habitat choice can be genetically variable, and so evolvable, within populations (Jaenike & Holt, 1991; Brown & Brown, 2000). Subsequently, habitat choice can be viewed as a mechanism that modulates - and likely moderates-selection on other phenotypic traits. Habitat choice may itself also be under (direct) selection. For example, as noted above, predation can result in changes in habitat use. Subsequently, differing selection pressures (as well as stochastic processes) can result in divergence in mean habitat preference among populations (Akcali & Porter, 2017; Satterfield & Johnson, 2020). Here we suggest multi-population, comparative studies offer an additional tool to investigate adaptive habitat choice that is currently under-exploited.

Scototaxis, or preference for dark versus light background environments, is an aspect of habitat choice that is also widely interpreted as reflecting shy-bold personality (Tudorache *et al*., 2013) or an ‘anxiety-like’ behavioural phenotype (Maximino *et al*., 2010b). Scototaxis assays present subjects with a choice between the safer (dark) background, and the riskier (light) background (Maximino *et al*., 2010b). On average, fish prefer dark over light backgrounds, although the strength of this preference differs among species (Maximino *et al*., 2007; Thompson *et al*., 2016; Jones *et al*., 2021). Additionally, evidence for among-individual variation in scototaxis was found in a captive population of guppies derived from the pet trade (De Russi *et al*., 2025).

The Trinidadian guppy provides a well-known model for studying the evolution of phenotypes (including behaviours) under predator-induced selection (Endler, 1984; Harris *et al*., 2010; Heckley *et al*., 2025). Across the mountainous Northern Range of Trinidad, waterfalls limit upstream movement of predators creating a pattern of high predation (downstream) and low predation (upstream) populations which is repeated across different rivers (Endler, 1984; Reznick & Bryga, 1996; Reznick *et al*., 2001). Predation regime has been shown to influence many guppy phenotypes, including behaviour (Endler, 1984; Schirmer *et al*., 2019; Whiting *et al*., 2022; Yong *et al*., 2022). Behavioural differences among wild guppy populations (and among captive lines) reflect a combination of environmental and genetic factors (Bleakley *et al*., 2006; Fischer *et al*., 2016). There is a general tendency for guppies from high predation sites to exhibit ‘bolder’ (i.e. more risk prone) behaviours on average (Harris *et al*., 2010). Experimental evidence suggests bold, exploratory individuals have higher survival in the presence of cichlid predators, indicating being bold in the face of predation may be adaptive (Smith & Blumstein, 2010). Within populations, behavioural traits associated with shy-bold variation (e.g. freezing, exploration in an open field trial) are typically repeatable (Burns, 2008; Houslay *et al*., 2018) and heritable (White *et al*., 2018; White & Wilson, 2018; Prentice *et al*., 2020; Houslay *et al*., 2022). Although much is known about guppy behaviour, the vast majority of prior studies of guppies have focused on adult fish, often motivated by investigating sexual behaviour and dimorphism (Harris *et al*., 2010; White *et al*., 2018; De Russi *et al*., 2025). Much less is known about patterns of variation within and among populations in early life. This is important as the strength of selection can vary across life stages (Camacho & Hendry, 2020). Crucially, the widely used high versus low predation categorisation of sites is based principally on the presence of large fish species that prey on adult guppies (e.g. the pike cichlid). Smaller, gape limited predators (notably *Rivulus hartii*) are present at all sites and may even impose greater mortality risk on juvenile guppies at nominally ‘low predation’ sites (Reznick *et al*., 1996).

Here we investigate scototaxis in juvenile guppies assayed in early life (up to 15 days post-birth). We use a common garden approach to test for evolutionary divergence in scototaxis among 10 captive populations of known wild origin. Using double hierarchical mixed models, we test for among-population variation in both mean scototaxis behaviour, and in levels of within-population variation. Among-population variation in a common environment can be attributed to genetic and therefore evolutionary divergence. However, this could reflect divergence via neutral as well as adaptive processes. To assess whether local adaptation to predation regime is a likely contributor to divergence, we assess the directional consistency of phenotypic differences between high and low predation populations within rivers. Simultaneously, we estimate behavioural variation among families within-populations. This allows us to characterise broad-sense heritable variation and so assess the adaptive potential within the populations.

## Methods

### Animal husbandry

We collected juveniles from 10 wild-type captive populations founded by wild fish collected from 6 rivers in Trinidad from stock tanks in the aquatic facility at the University of Exeter’s Cornwall campus (UK) over a 6-month period from August 2021 to April 2022. To obtain juvenile guppies from known families, we opportunistically isolated visibly pregnant females. Paired high predation (HP) and low predation (LP) populations were available for four rivers (Aripo, Guanapo, Quare, and Yarra). We also paired a high predation population from the Marianne, and a low predation population from the adjacent Paria. Despite being on different rivers, these populations have been shown to be closely related and have extensive gene flow (Willing *et al*., 2010; Blondel *et al*., 2019). Eight of the populations were founded from collections made in 2017 at sites described in (Yong *et al*., 2022) (Table 1). A ninth population (ALP) was founded in 2020 by combining fish descended from two distinct (though genetically similar) low predation sites in the Aripo sampled in 2017 (Yong *et al*., 2022). This was to reduce husbandry workload during covid-related facility access restriction. A final population (Guanapo low predation, GLP) was acquired from a stock descended from an earlier wild collection in 2012 (D. Croft, personal communication).

**Table 1:** Description of guppy populations tested including wild collection sites for origin stock and sample sizes used.

| River | Predation regime | Latitude | Code as used in Yong et al. 2021 | Longitude | Drainage | Wild collection year | Number of families tested | Number of juveniles tested |
| --- | --- | --- | --- | --- | --- | --- | --- | --- |
| Aripo | High | 10° 39' 02" | AH | 61° 13' 26" | Caroni | 2017 | 24 | 149 |
|  | Low | 10° 41' 08" | AL * | 61° 13' 56" | Caroni | 2017 | 15 | 58 |
|  |  | 10° 41' 10" | AL2 * | 61° 13' 30" | Caroni | 2017 |  |  |
| Guanapo | High | 10° 39' 28" | GH | 61° 15' 13" | Caroni | 2017 | 8 | 35 |
| Guanapo | Low | NA | GL | NA | Caroni | 2012 | 19 | 82 |
| Marianne | High | 10° 45' 54" | MH | 61° 18' 15" | Northern | 2017 | 11 | 31 |
| Paria | Low | 10° 44' 57" | PL | 61° 15' 54" | Northern | 2017 | 23 | 88 |
| Quare | High | 10° 39' 53" | QH | 61° 11' 35" | Oropuche | 2017 | 7 | 27 |
| Quare | Low | 10° 40' 33" | QL | 61° 11' 49" | Oropuche | 2017 | 10 | 45 |
| Yarra | High | 10° 47' 23" | YH | 61° 21' 12" | Northern | 2017 | 10 | 40 |
| Yarra | Low | 10° 44' 25" | YL | 61° 19' 17" | Northern | 2017 | 8 | 31 |

\* During covid-19 lab closures lab stocks of the Aripo low predation populations referred to as AL and AL2 in Young *et al.* 2021 were combined to ease husbandry requirements.

Visibly pregnant females were individually transferred from population-specific stock tanks (30L) into 2.9L breeding tanks (22cm × 8.5cm × 15cm) with an independent recirculating water supply (supplied by Aquaneering). Breeding tanks were checked daily for fry, and females were returned to stock tanks after giving birth to avoid risk of fry predation. Offspring then remained in family groups, fed twice a day with commercial flake food and laboratory-prepared *Artemia salina* nauplii, until behavioural testing. After testing they were returned to designated stock tanks. Females were isolated on an ad hoc basis and since time from isolation to birth was variable, families became available for testing asynchronously. We therefore conducted behavioural testing in batches (at up to 2-week intervals) depending on availability of juveniles. As a result, the age at testing varied among families from 1-15 days (with a mean among families of 6.74 days). In total we collected 586 juveniles from 135 families. The number of juveniles tested per family ranged from 1 to 33 with a mean of 4.43, whilst the number of families per population ranged from 7 to 24 with a mean of 13.5 (see Table 1).

### Behavioural testing

Guppies were individually tested in a scototaxis assay when aged between 1 and 15 days old, with all members of a family tested on the same day. Testing was conducted in a cubic trial tank (10 × 10 × 10 cm) filled to a depth of 5cm using water from the breeding tank in which the focal individual had been housed. Water temperature was recorded for the majority of trials, and ranged from 21-24°C with a mean of 23.1 °C. However, due to experimenter oversight this parameter was not recorded for 167 of the 586 trials conducted. The trial tank arena was comprised of two equally sized but visually distinct sides (Figure 1). Floor and walls were black on the dark side and pale grey on the light side. Indirect lighting was provided from a nearby lamp. Fish were introduced to the centre of the arena by being transferred, using a turkey baster, from their breeding tank into a centrally positioned opaque tube (that matched the wall colours of the arena (Figure 1). After a 30s acclimation period, the tube was lifted and the individual’s location was tracked for 300 seconds using a Sunkwang C160 video camera with 6–60 mm manual focus lens suspended above the tank using the tracking program Viewer II (https://www.biobserve.com). We defined a single trait of ‘preference’ as the difference (in seconds) between time spent on the light and dark sides of the arena, with positive values denoting more time spent on the light side. While the tracking software could readily locate which side of the arena each fish was on, a lack of contrast against the dark background precluded reliable extraction of additional traits (e.g. activity as measured by total track length swum).

**Figure 1:**
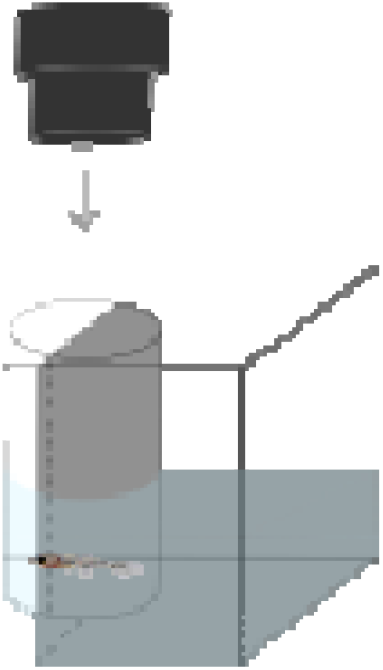
Apparatus used to assay light-dark preference (scototaxis) response of juvenile guppies. The tank is visually separated into two halves with light/dark grey walls and floor. Guppy juveniles were introduced into a centrally placed tube to acclimatise before being released. Movement was tracked using an overhead camera.

### Data analysis

We plotted data by predation regime and population to assess whether any patterns were visually apparent before modelling preference using two mixed effect models fitted with R package brms (v. 2.22.0)(Bürkner, 2017), which is based on the stan software (Carpenter *et al*., 2017), in R (v. 4.4.3)(R Core Team, 2025). We assume Gaussian errors and acknowledge this assumption is inevitably violated by our defined response variable (*preference*) which is bounded between -300 and + 300 seconds. We note that observations could be treated as proportions (e.g. of time spent in light) but that they do not arise from a binomial process. Observed preference scores show a bimodal distribution within each predation category (Figure 2A), and this is also apparent within at least some of the individual populations (e.g. Aripo HP, Figure 2B). Since bimodal distributions are not readily approximated by a beta distribution we opted to use (Gaussian) linear mixed models (LMM) and inference and parameter estimation are known to be generally robust to deviations from normality (Schielzeth *et al*., 2020). In practice inspection of model residuals suggested that the assumption of a Gaussian error structure was not unreasonable.

**Figure 2:**
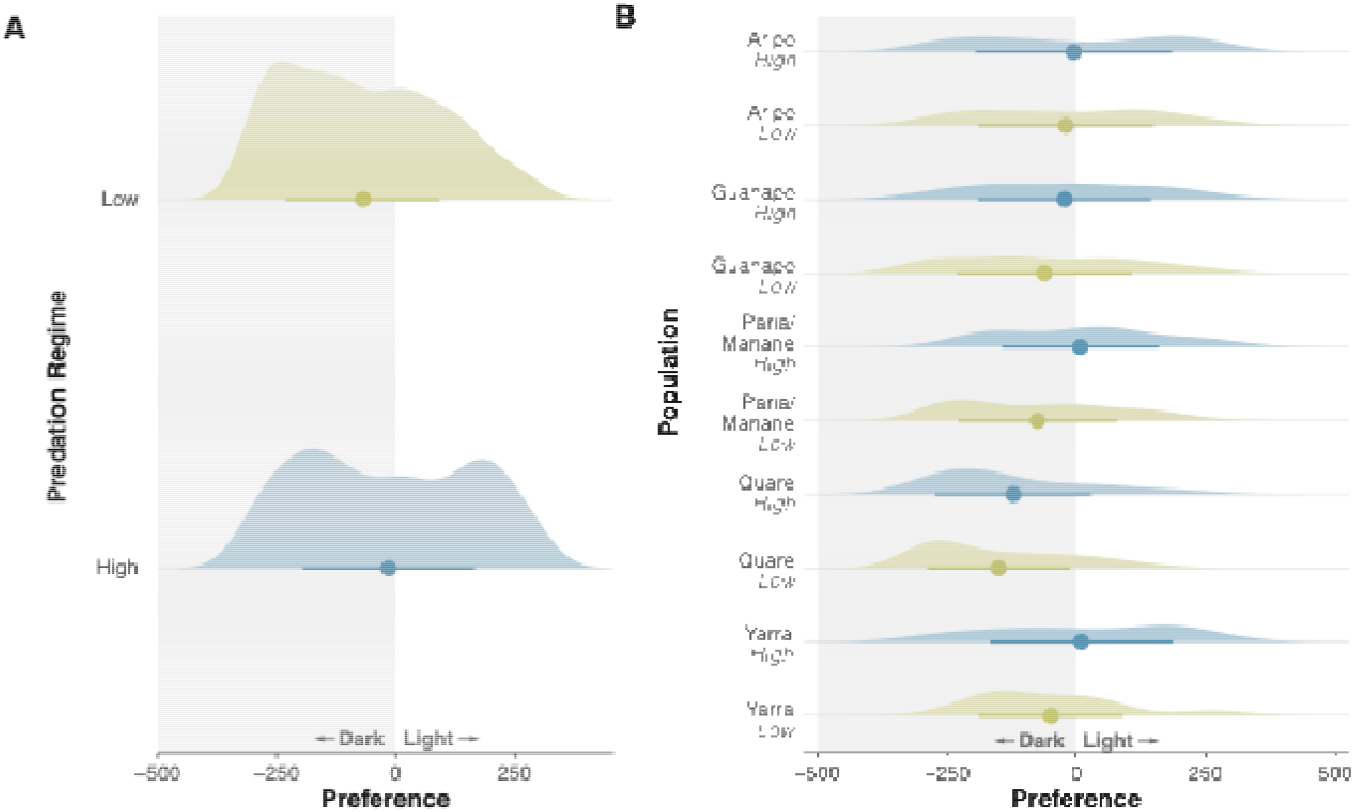
Observed preference distributions for juvenile guppies shown by A) different ancestral predation regimes and B) different ancestral populations. Points indicate mean preference whilst the bar indicates standard deviation from the mean. Low predation populations are shaded in green whilst high predation populations are blue.

First, we used a linear mixed model (LMM) to jointly estimate variance in *preference* among populations and among families within populations. We refer to this as the ‘*mean model’* since we used it to characterise variation in mean behaviour among populations and among families. The *mean model* included fixed effects of age and temperature as linear covariates (standardised to a zero mean). Missing temperature records were assigned a value of zero (i.e. the mean of observed records). Within-family trialling order was also included as a covariate to account for any cumulative disturbance caused by serially removing fish from their home breeding tank (Prentice *et al*., 2020). Note that we did not include predation regime as a fixed effect because we wanted to estimate (total) among-population variance using the random effect structure (as opposed to among-population variance not explained by predation regime). Both population and family were included as random effects. We estimated adjusted repeatabilities, or intra-class correlations (ICC), at both family and population level, by expressing the corresponding variance components as a proportion of the total phenotypic variance (V_P_) conditional on the fixed effects (Cleasby, Nakagawa and Schielzeth, 2015). The total phenotypic variance (V_P_) was estimated as the sum of the variance explained by population (V_Pop_), family (V_Fam_) and the residual variance (V_Res_). We used the posterior means of each variance component as our point estimates with 95% credible intervals estimated using HPDinterval() from brms. Broad sense heritability was estimated, under an assumption of full-sibling families, using H^2^= 2V_fam_/2V_fam_ + V_pop_ + V_res_ (Falconer & Mackay, 1960). Broad sense heritability estimates do not distinguish between total genetic variance and the additive genetic component that determines transmission of traits from parents to offspring. As such, to the extent that non-additive genetic effects are present, H^2^ will overestimate a trait’s potential to respond to directional selection (Visscher *et al*., 2008). We also reiterate that our estimate is assumption laden as true paternities are unknown (i.e. there may be mixed paternity within some maternal sibships and/or shared paternity across some maternal sibships within populations).

Our second model extended the above analysis using a double hierarchical mixed model (DHGLM). This allowed us to; (i) test for differences in the variance in preference among-populations and families; and (ii) estimate any relationship between the mean *preference* and the within group variance at these two hierarchical levels of behavioural variation. We refer to this as the ‘*variance model*’. The variance model of *preference* included the same fixed and random effects detailed above, as well as random effects of population and family groups on residual variation modelled on a log scale. We also estimated a covariance between mean *preference* (adjusted for fixed effects) and intra group variance of *preference* at population and family levels. While noting such relationships can arise from distributional considerations as well as biological ones (see Discussion), this allowed us to ask whether groups (i.e., populations, families) with high (or low) mean behaviours also tend to contain high (or low) levels of behavioural variation.

We used default weakly informative priors in brms. The intercept and random effects were fitted with Student’s t priors with three degrees of freedom, whilst the fixed effects were fitted with flat priors. We ran four chains with a warmup of 1000 iterations and 2000 post warmup iterations with a thinning interval of one. Therefore, all estimated model coefficients and credible intervals were based on 4000 posterior samples. All parameters had Ř < 1.00, as well as both bulk and tail-estimated sample sizes > 1000, indicating satisfactory convergence. This was additionally confirmed through visual inspection of model performance (Supplementary Figures 1 and 2). We additionally validated our mean model by comparing parameter estimates to the results of a REML (restricted maximum likelihood) fit using lmer4 (v. 1.1.37, Bates *et al*. 2015) and lmerTest (v. 3.1.3, Kuznetsova *et al*., 2017).

## Results

Across all individuals, mean observed preference was negative (mean ± SE = -43.29±7.13) indicating juvenile guppies tended to prefer the dark side of the arena on average. Guppies from low predation (LP) populations have a stronger preference for the dark than HP populations (mean_LP_ ±SE =-69.4 ± 9.24, mean_HP_ ± SE= -15.1 ± 10.7, t=3.84, df= 564.59, P < 0.001) (Figure 2A). Visual inspection of the raw data by population confirmed this pattern holds qualitatively within the four rivers with directly comparable high predation and low predation populations (Figure 2B). Thus, mean preference is lower (i.e. stronger preference for the dark) in the LP populations across all rivers.

More formally, our *mean mode*l yielded a negative intercept estimate supporting the conclusion of a slight preference for the dark on average over all fish (intercept = -39.26, 95% CI= -83.94, 3.34; Table 2). This estimate is biologically interpretable as the expected preference for a fish of average age, tested first in its group, and at an average temperature. Though not directly relevant to *a priori* hypotheses, our *mean model* also indicates a positive effect of temperature on preference for the light, while effects of age and order are negative (though 95% CI spanned zero for the order effect). ICC estimates from this model indicate that, conditional on fixed effects, population and family explain 11 and 12%, respectively, of variation in preference (Table 3). Note that variance component estimates are constrained to be positive, and therefore Bayesian credible intervals on the components (or derived ICC) cannot be used for inferring significance in a frequentist sense. Here the lower 95% credible interval for the population-level ICC is close to zero, but the posterior mean is nonetheless distinct from zero (Supplementary Figure 3). The posterior of the family-level ICC is clearly distinct from zero. We therefore consider both among-population, and among-family within population variance to be supported. For comparison, refitting the *mean model* in a frequentist framework using LMM and testing random effects by likelihood ratio test, yielded similar ICC estimates and confirmed both random effects were significant at P<0.05 (Supplementary Table 1). The model yielded a broad sense heritability estimate of H^2^=0.20 (CI=0.08, 0.33), though we reiterate that this estimate makes the unverified assumption that our data set comprises full sibling families only.

**Table 2:** Estimated fixed and random effects from the *mean model* of preference. Preference was measured as the difference between time in the light and dark, such that the negative intercept indicates preference for dark. Random intercept variances are shown on a standard deviation scale (σ). Values in square brackets denote 95% credible intervals.

| Model term | Estimate |
| --- | --- |
| *Fixed Effects* |  |
| Intercept | -39.26 [-83.94, 3.34] |
| Age (Standardised) | -6.26 [-10.03, -2.37] |
| Order | -4.26 [-9.65, 1.11] |
| Temperature (Standardised) | 26.93 [10.90, 43.23] |
| *Random effects* |  |
| $\sigma_{\text{Population}}$ | 55.94 [22.12, 106.78] |
| $\sigma_{\text{Family}}$ | 58.57 [37.06, 80.49] |

**Table 3:**
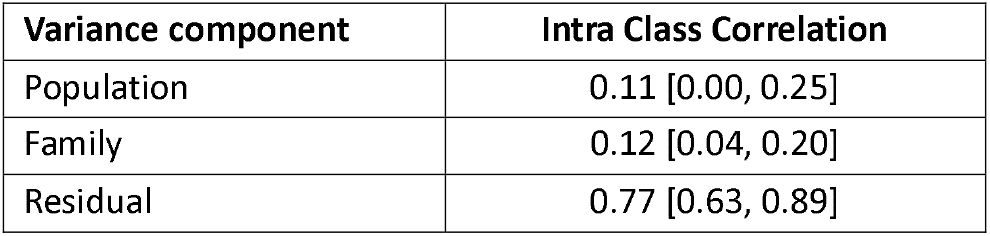
Estimated intra-class correlations (ICC) for among-population, among-family and residual variance components from the *mean model* of preference. Values in square brackets denote 95% credible intervals.

| Variance component | Intra Class Correlation |
| --- | --- |
| Population | 0.11 [0.00, 0.25] |
| Family | 0.12 [0.04, 0.20] |
| Residual | 0.77 [0.63, 0.89] |

As expected, the *variance model* yielded very similar estimates of fixed effects and random effect variances for the mean population and family effects, so we do not detail these further here (but see Tables 2 and 4 for all parameter estimates). The posterior distribution for among-family variance of within-family variation is distinct from zero (shown as the standard deviation of family specific standard deviations on a log-scale in Supplementary Figure 4). The same is true at the population level with the DHGLM indicating that some populations harbour more variation than others. Among populations, high predation sites tend to be less variable (i.e. contain more variation around the population mean) than their low predation counterparts from the same river (Figure 3A). This was the case for 4 of our 5 paired comparisons, the exception being in the Guanapo river, where the low predation population is less variable. There was a moderate - but highly uncertain - positive correlation between population mean preference and intra-population variation in preference (r=0.55 CI=-0.59, 0.99; Table 4). Thus, the qualitative pattern is that populations with stronger mean preference for the light (or weaker mean preference for the dark) tend to be more variable but this relationship is not statistically supported.

**Table 4:** Estimated fixed and random effects from the *variance model* of preference fitted using a Bayesian double hierarchical mixed effect model. Fixed effects were as specified for the *mean model*. Random intercept variances are shown on the standard deviation scale (σ), as are Population and Family effects on residual variance (on the log scale, ω) and correlations between group level effects on mean and variance (r). Values in square brackets denote 95% credible intervals.

| Model term | Estimate |
| --- | --- |
| *Fixed Effects* |  |
| Intercept | -40.38 [-86.85, 2.31] |
| Age (Standardised) | -6.56 [-10.44, -2.76] |
| Order | -4.21 [-9.71, 1.06] |
| Temperature (Standardised) | 26.03 [9.70, 42.77] |
| *Random effects* |  |
| $\sigma_{\text{Population}}$ | 55.22 [22.91, 104.83] |
| $\omega_{\text{Population}}$ | 0.10 [0.01, 0.24] |
| $r_{\text{Population}}$ | 0.55 [-0.59, 0.99] |
| $\sigma_{\text{Family}}$ | 63.37 [43.71, 83.16] |
| $\omega_{\text{Family}}$ | 0.11 [0.01, 0.22] |
| $r_{\text{Family}}$ | 0.72 [-0.26, 0.99] |

**Figure 3:**
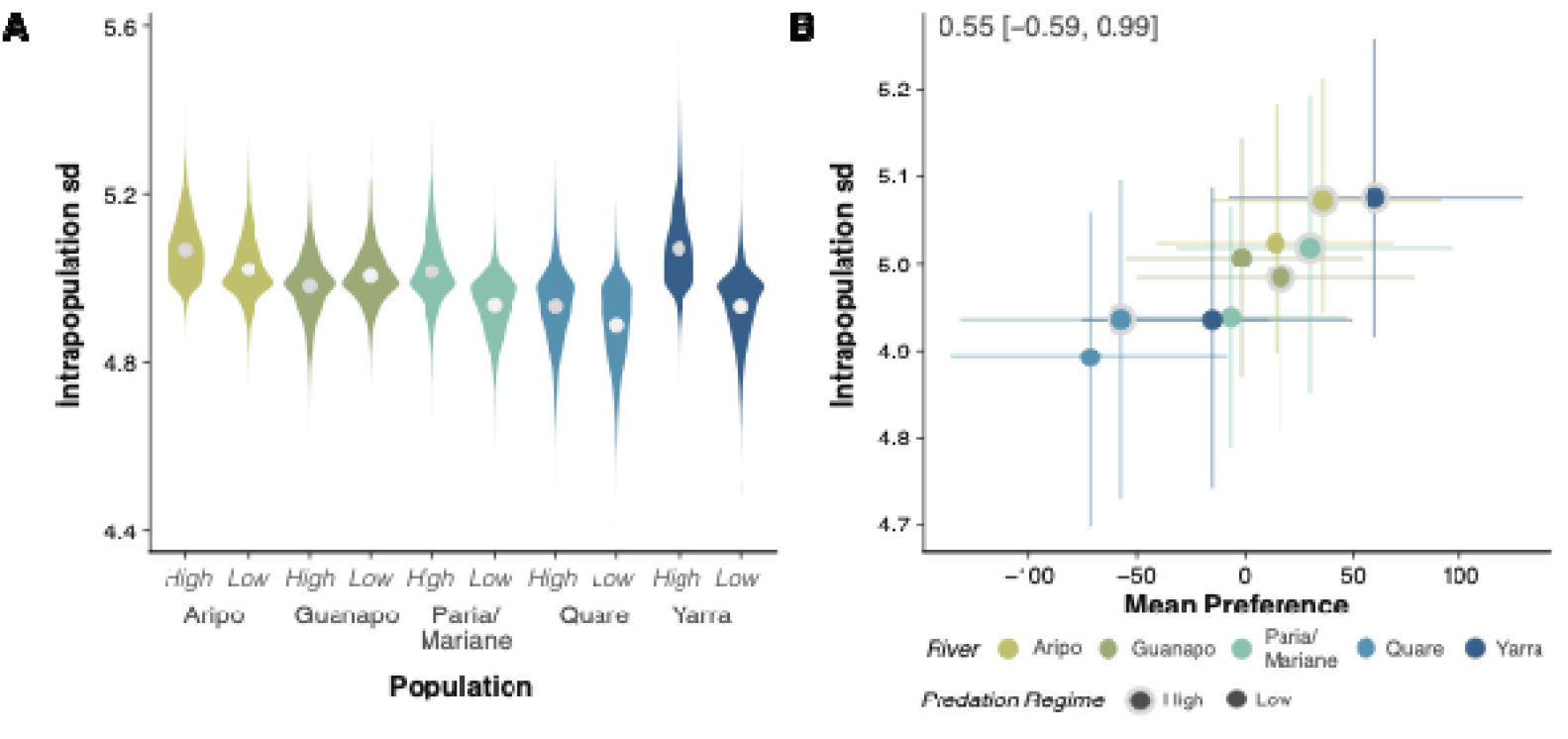
A) Estimated intrapopulation variance of each population, from the *variance model* of preference, shown here as the standard deviation (sd). Violin plots indicate the distribution for the population-specific posteriors, with the posterior mean denoted by a point. Populations are grouped by river and predation regime to facilitate visual comparison. B) The relationship between model predictions of the mean preference and intrapopulation variance (shown as standard deviation) among populations. The colour of the central point indicates the river of origin, and the presence of a grey outline denotes high predation populations. The lines show 95% credible intervals.

The intra-family variance also differs among families within populations (0.14, CI=0.01, 0.28; Table 4, Supplementary Figure 5A). Unsurprisingly, since family effects are estimated conditional on other terms in the model, differences among families are not aligned with river or ancestral predation regime (both of which are captured by the population effect: Supplementary Figure 5A). There is a strong positive correlation between family mean and intra-family variation (r=0.77 CI=-0.17, 1.00; Table 4); families with stronger mean preference for the light had more variability in response, a pattern seen within each population (Supplementary Figure 5B). While the posterior mode for this correlation was clearly distinct from zero (Supplementary Figure 4) uncertainty was again high, with the 95% credible interval spanning zero (86.5% of the posterior was >0).

## Discussion

Here we used a scototaxis assay to characterise variation in a habitat choice trait, preference for a light vs dark background, both within- and among-populations of juvenile guppies raised under common garden conditions. We find a weak preference for the dark microhabitat in juvenile guppies on average, which aligns with findings in other small fishes (Maximino *et al*., 2007, 2010a; Wagle *et al*., 2017; Pecunioso *et al*., 2024), and adult guppies (De Russi *et al*., 2025). Additionally, we show mean preference varies among families, consistent with this being a heritable trait which could evolve under selection. We also find mean preference varies among populations which, under the common garden conditions used, provides evidence for evolutionary divergence. This is consistent with the hypothesis that adaptation to local predation regime has contributed to this divergence. Juvenile guppies descended from high predation populations tend to be bolder, with a lower mean preference for the dark microhabitat, than those from low predation environments in the same river. Alongside differences in mean behaviour, we also find that families and populations vary in how much behavioural variation they contain. At both levels, weaker mean preference for the dark is associated with the presence of more intra-group variance, indicating that bolder populations and families are more variable in their scototaxis response.

Here, we find strong support for variation in mean preference among families, within populations. Therefore, there is support for broad sense heritability, and thus evolutionary potential of juvenile scototaxis. However, it is important to highlight some limitations and partial caveats to this conclusion. Maternal effects on open-field behaviours have been shown in juvenile guppies from the same Aripo high-predation site used here (White & Wilson, 2018) and therefore they may contribute to the among-family variation in scototaxis shown here. However, earlier work also concluded maternal effects were partially explained by both female size and brood size (White & Wilson, 2018), both traits which are expected to be heritable (Whiting *et al*., 2022). Thus, at least some maternal effects arise from genetic (among-mother) variation and so contribute to ‘total heritability’ of offspring traits (Willham, 1972; Wilson *et al*., 2005; Wolf & Wade, 2016). Here we lack data on maternal size, but brood size (which is positively correlated to female size in guppies; Reznick 1983) is known. Refitting the model with brood size yielded some support, with a positive effect of brood size (and 96% of the posterior > 0) and reduced family variance by 8.6 %, relative to our *mean model* estimate (Supplementary table 2). This is consistent with the hypothesis that brood size is a source of pre-natal maternal effects on juvenile behaviour. However, brood size also determines post-natal housing density in the short period until behavioural testing (i.e. from 1-14 days), which could equally be viewed as a common-environment effect. More generally, we tried to reduce the potential for common-environment effects by using a recirculating water supply but lacked the facility space to split families across multiple tanks. We therefore cannot completely rule out a contribution of post-natal common environment (i.e. tank) effects to among-family variation.

After finding evidence for heritable variation in light/dark preference, we then investigated whether there was divergence among populations and, if so, whether this was consistent with local adaptation to predation regime. We found support for population level evolutionary divergence in scototaxis response, with 11% of variance explained by population. As populations have been maintained in a ‘common garden’, we interpret this as quantitative genetic variance among-populations arising from an unknown combination of neutral and adaptive processes (i.e. divergence by drift under restricted gene flow and local adaptation). Juveniles from low predation populations showed a significantly stronger average preference for the dark overall, a pattern which was qualitatively repeated in four out of five within-river comparisons between LP and HP populations. This pattern is suggestive of high predation environments selecting for bolder phenotypes. It also concurs with prior work on adult guppies showing fish from high predation populations have shorter emergence and hesitancy times than those from low predation populations (Harris *et al*., 2010).

We currently do not understand the fitness consequences of variation in scototaxis response in wild populations, so any adaptive interpretation is speculative. However, incorporating predation risk into the marginal value theorem suggests the optimal level of boldness should increase with risk (Calcagno *et al*., 2025); meaning when predation risk is high, individuals must be bold to gain resources.

Notably, it is empirically challenging to separate a behavioural response to risk from the perception of risk itself. An alternative hypothesis is, therefore, that the level of risk presented by our experimental paradigm is being perceived as greater on average by fish from populations with low predation ancestry. Regardless, we see evolutionary divergence of juvenile scototaxis among populations and patterns are qualitatively consistent with adaptation to predator-mediated selection contributing to this.

The intensity of selection is thought to vary across age classes and is considered to be especially important in fishes, where many predators are gape-limited (Arendt & Reznick, 2005). Subsequently, predation pressure is likely highest in the youngest and smallest cohort. However, our mean observed preference score of -43.3s corresponds to 57% of time on the dark side, exactly matching the average recently reported in adult guppies from a domestic population (De Russi *et al*., 2025). Thus, to the extent we can directly compare these studies, average scototaxis response appears very similar in juvenile and adult guppies. If predator driven selection was greater in juveniles than adults, we would expect preference for the dark to be greater in juveniles. However, as we have shown in this study, the mean light/dark preference varies among populations, meaning that a valid comparison would also require examining the scototaxis response in adults from each population used here.

As well as exploring differences in mean scototaxis at family and population levels, we also modelled differences in the amount of variation contained within these grouping factors. Families using the light side more on average contain more behavioural variation, a pattern repeated (albeit with less statistical support) when comparing among populations. Interpretation of these patterns is non-trivial as there are multiple possible biological drivers, and because mean-variance relationships can sometimes arise from distributional properties of the focal trait (Tatliyer *et al*., 2019). Prior applications of DHGLM have mostly explored intraÖindividual variability, but such analyses require repeated observations of known individuals, which was not feasible in this study (Cleasby *et al*., 2015; Jolles *et al*., 2019). Hence, we cannot yet say whether families with higher intra-family variance contain more personality variation among-siblings, or more within-individual variation, or both. Nonetheless, if we accept family level differences in mean behaviour as putatively indicative of heritable genetic variation, then applying the same logic to intra-family variance suggests the presence of heritable variation for variability of the scototaxis response. This implies a role for genotype-by-environment (GxE) interaction (Rönnegård *et al*., 2010; Prentice *et al*., 2020). This aligns with the conclusions of Prentice *et al*. (2020) who detected GxE for a behavioural stress response trait in adult guppies from the Aripo high predation population by estimating additive genetic variance for individual ‘predictability’. Interestingly, in that study, genotypes (and individuals) with bolder mean phenotypes were more predictable (i.e. less variable). Here we find the converse at the family-level; putatively bold families that spend more time in the light on average are less predictable (i.e., contain higher intra-family variance). While the biological significance of this remains unclear, a meta-analysis shows that correlations between behavioural means and predictabilities are abundant but vary in direction (Horváth *et al*., 2023). We note that mean-variance correlations can sometimes follow inevitably from the trait distribution (i.e., a correlation of zero is not necessarily an appropriate null model), we do not think this explains the current results. Specifically, we analysed *preference* under Gaussian assumptions but, if viewed as a proportion we might expect highest variance in families with mean *preference* of zero (i.e. equal probability of being on light or dark sides) which is opposite to the pattern seen.

At the population level, our *variance* model revealed a trend towards high predation populations containing more variation than low predation populations from the same river. Acknowledging the limitations imposed by the low number of available high predation vs low predation population pairs, there are adaptive hypotheses that predict this pattern. First, behavioural unpredictability could itself be an adaptation to predation risk if highly consistent, stereotypical responses to risk increase the likelihood of being caught. In hermit crabs (*Pagurus bernhardus*), emergence time becomes less predictable under elevated predation risk (Briffa, 2013), while interactions with humans reduce predictability of movement patterns in brown bears (*Ursus arctos*) (Hertel *et al*., 2021). In these examples, changes in predictability arise from plasticity, occurring within individuals according to perceived risk. However, local adaptation could equally lead to divergence among-populations with different selection regimes, provided genetic variance is present. Second, if shy-bold variation among-individuals is maintained by predator-mediated risk-reward trade-offs, as has been widely proposed (Moiron *et al*., 2020), then we would predict more behavioural variation in populations with greater predation risk. However, there are also plausible neutral models to consider; upstream, low predation guppy populations have consistently lower effective population size (Fraser *et al*., 2015), such that they lose genetic, and so phenotypic variation, more rapidly through drift. While testing these hypotheses is beyond the scope of the present study, it highlights how application of DHGLM to multi-population data sets could facilitate comparative approaches to the studies of animal personality variation (White *et al*., 2020; Dalos *et al*., 2022).

In conclusion, we show here that scototaxis is likely to be a heritable trait in juvenile guppy populations, and that patterns of evolutionary divergence among-populations are consistent with local adaptation to predation regime having shaped both the mean and variance of population-specific behavioural distributions. Our work suggests that lower predation populations have both a stronger preference for the dark and are less variable in their scototaxis response than high predation populations from the same river. Taken together, our results support the premise that there is variation in habitat choice traits linked to shy-bold phenotypes, which could be acted upon by natural selection. We also hope that our results illustrate the value of employing multi-population comparisons under common-garden conditions for understanding the evolution of behavioural processes under natural selection, a framework which has not been widely adopted in behavioural studies.

## Acknowledgements

We thank Tom Black and the animal care staff in the aquatic facility for assistance with animal husbandry.

