## Supplementary Materials for "Adaptive potential and evolutionary divergence of scototaxis in juvenile Trinidadian Guppies" (from another version of this preprint: version 1, 2026-05-05)

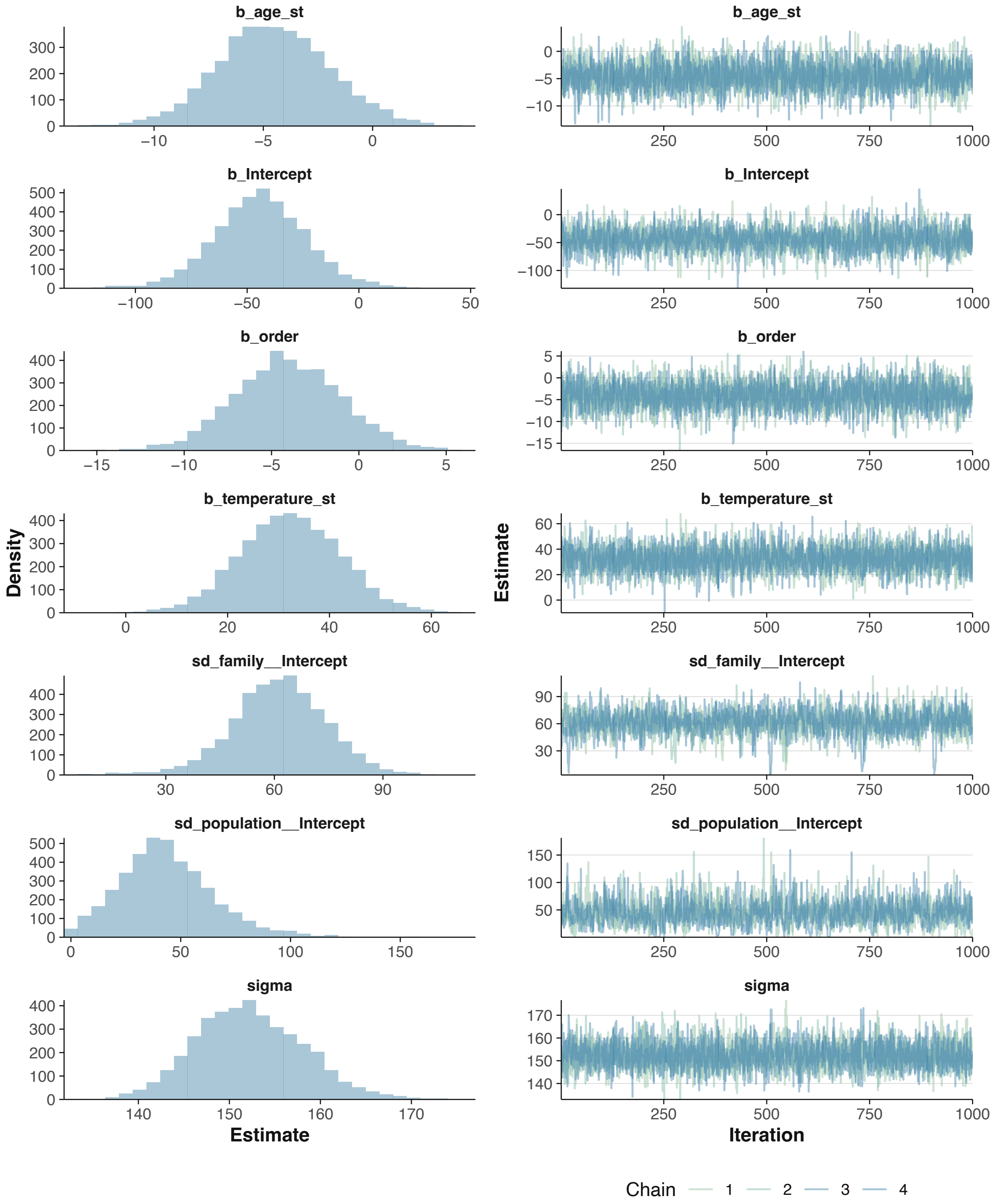

**Supplementary Figure 1:** The density of the estimates and convergence of chains of the *mean model* of preference, a Bayesian mixed effect model exploring the light/dark preference in juvenile guppies.

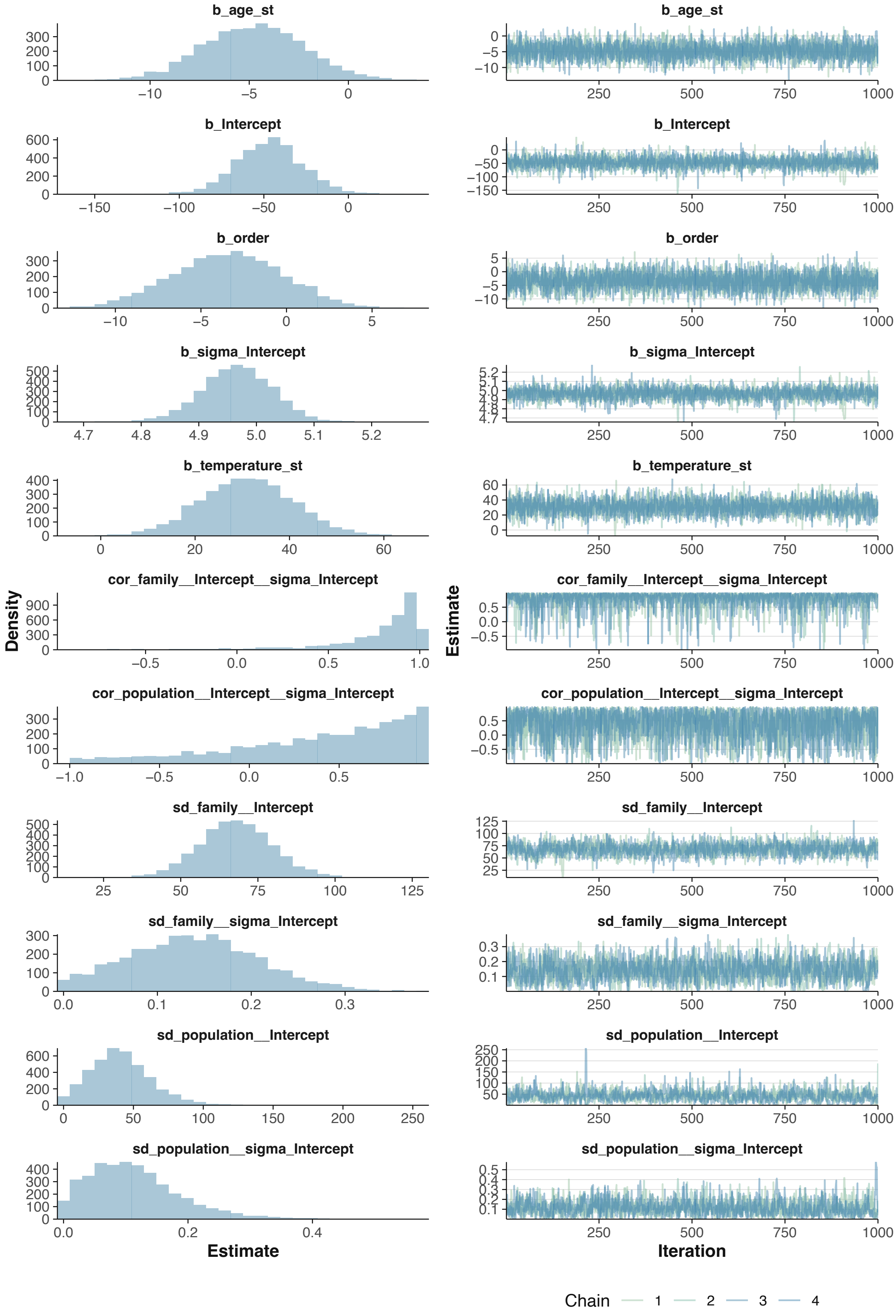

**Supplementary Figure 2:** The density of the estimates and convergence of chains of the *variance model* of preference, a Bayesian double hierarchical mixed effect model exploring the light/dark preference of juvenile guppies.

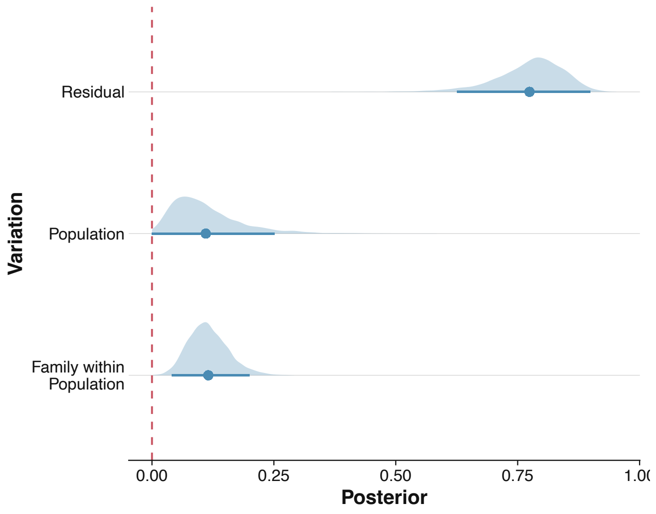

**Supplementary Figure 3:** The distribution of posteriors of the variance (Intra Class Correlation) attributed to among-population, among-family and residual variation estimated from a Bayesian fixed effect model of light-dark preference. The dark point indicates the mean posterior value whilst the dark blue line shows the 95% confidence intervals.

**Supplementary Table 1:** Estimated fixed and random effects from the *mean model* of preference (difference between time in the light and dark) refitted using restricted maximum likelihood (REML) in lme4. Random effect variances are shown on the standard deviation scale (σ). Also shown are F-tests of fixed effects with denominator degrees of freedom estimated using Satterthwaite's method, and likelihood ratio tests of random intercepts. For testing each random effect, we assume the test statistic is distributed as a 50:50 mix of $\chi^2_0$ and $\chi^2_1$ (following Self and Liang (1987)).

| **Model term** | **Estimate (SE)** | **Statistical inference** | | |
|---|---|---|---|---|
| *Fixed effects* | | F | DF | P |
| Intercept | -39.73 (18.88) | 5.36 | 1, 14 | 0.036 |
| Age (Standardised) | -4.45 (2.51) | 3.15 | 1, 95.2 | 0.079 |
| Order | -4.23 (2.97) | 2.03 | 1, 358.4 | 0.155 |
| Temperature (Standardised) | 32.35 (9.77) | 10.97 | 1, 174.7 | 0.001 |
| *Random effects* | | $\chi^2_{0,1}$ | P | |
| $\sigma_{Population}$ | 39.60 | 3.46 | 0.031 | |
| $\sigma_{Family}$ | 60.85 | 12.51 | <0.001 | |

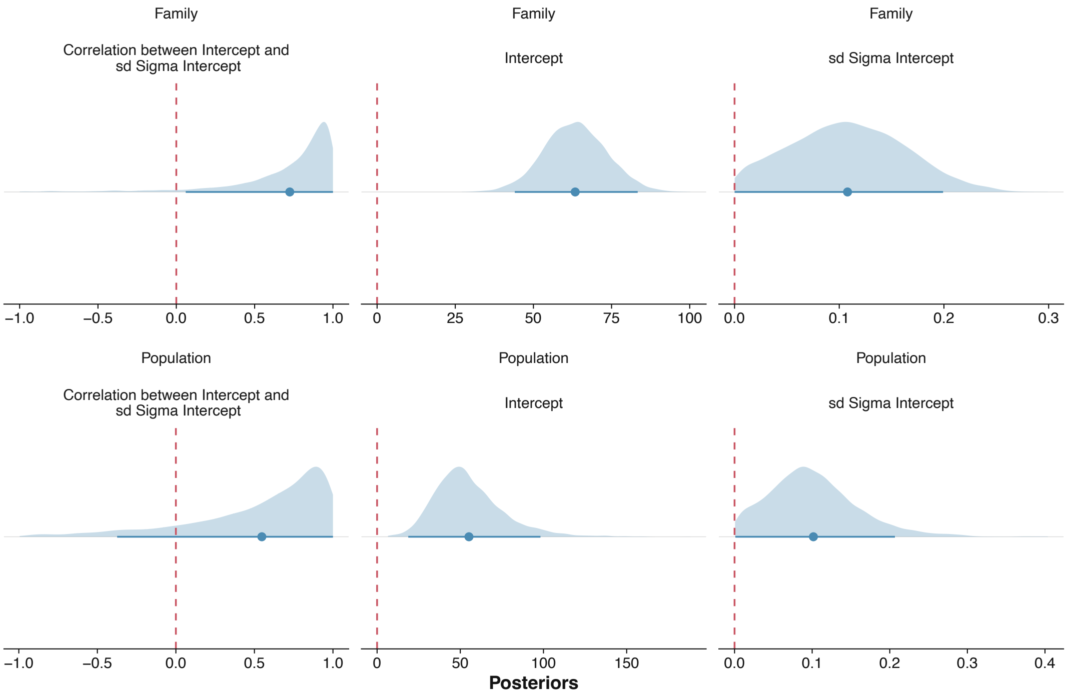

**Supplementary Figure 4:** Posterior distributions of parameters from the Bayesian double-hierarchical fixed-effects *variance model* of preference. Dark points indicate posterior means, and dark blue lines represent the 95% credible intervals.

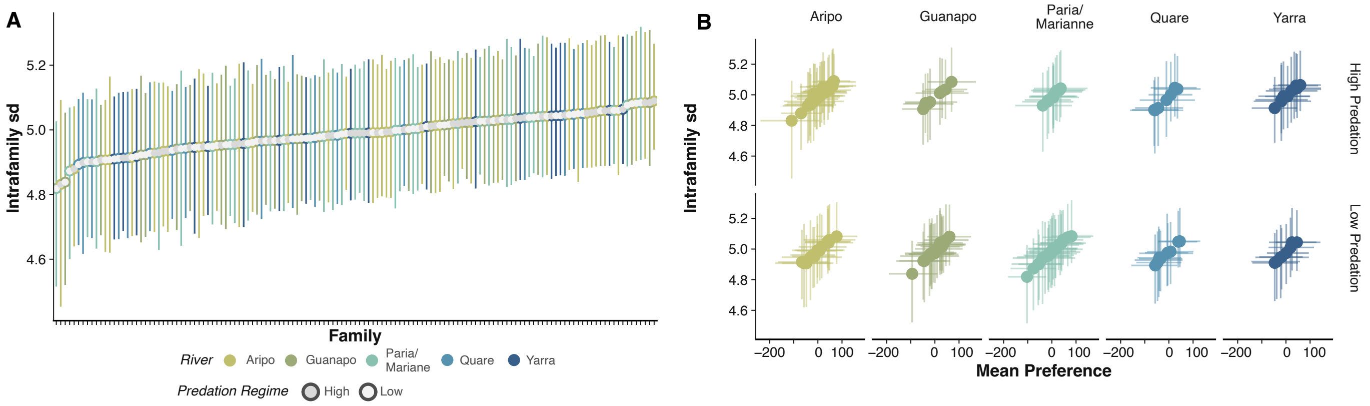

**Supplementary Figure 5:** A) Intra-family variance in light/dark preference among families, within populations, estimated from the *variance model* of preference, a Bayesian double hierarchical mixed effect model. Variance here is shown as sd on the log scale. The mean intra-family variance is denoted by a point whilst the range lines indicate the upper and lower 95 % confidence intervals. Families have been sorted by mean intrafamily variance, to highlight the lack of clustering across rivers or predation regimes. River and predation regime are indicated by the colour of the outline of the point range and the internal colour respectively. B) The correlation between mean preference per family and intrafamily variance among families, computed within the model. Families have been separated by both predation regime and by river of origin. The lines show the 95% confidence intervals.

**Supplemental Table 1:** Estimated fixed and random effects from the *mean model* of preference refitted with Brood Size included as an additional fixed covariate. Random intercept variances are shown on a standard deviation scale (σ). Values in square brackets denote 95% credible intervals.

| **Model term** | **Estimate** |
|---|---|
| *Fixed Effects* | |
| Intercept | -53.00 [-99.35, -7.77] |
| Age (Standardised) | -6.94 [-10.64, -3.11] |
| Order | -6.29 [-11.81, -0.58] |
| Temperature (Standardised) | 25.54 [9.52, 41.74] |
| Brood Size | 4.05 [-0.40, 8.37] |
| *Random effects* | |
| $\sigma_{Population}$ | 55.94 [22.12, 106.78] |
| $\sigma_{Family}$ | 58.57 [37.06, 80.49] |
